# Effect of Soybean and Seaweed-based Diets on Growth Performance, Feed Utilization, and Gut Microbiota of Tilapia: A Systematic Review and Meta-analysis

**DOI:** 10.1101/2023.10.20.563235

**Authors:** Leonildo dos Anjo Viagem, Jean Nepomuscene Hakizimana, Cyrus Rumisha, Brunno da Silva Cerozi, Gerald Misinzo

**Affiliations:** SACIDS Africa Centre of Excellence for Infectious Diseases, SACIDS Foundation for One Health, Sokoine University of Agriculture, PO Box 3297, Morogoro, Tanzania; Department of Animal, Aquaculture and Range Sciences, College of Agriculture, Sokoine University of Agriculture, PO Box 3004, Morogoro, Tanzania; Department of Food and Agricultural Sciences, Rovuma University, Cabo Delgado, Mozambique; Department of Animal Science, College of Agriculture, University of São Paulo, Avenida Padua Dias, 11, PO Box 9, Piracicaba, São Paulo, Brazil; Department of Veterinary Microbiology, Parasitology and Biotechnology, College of Veterinary Medicine and Biomedical Sciences, Sokoine University of Agriculture, PO Box 3019, Morogoro, Tanzania

**Keywords:** soybean, seaweed, diet, gut microbiota, tilapia, fish

## Abstract

Tilapia, a significant aquaculture species globally, relies heavily on feed for its production. While numerous studies have investigated the impact of soybean and seaweed-based diets on tilapia, a comprehensive understanding remains elusive. This review aimed at evaluating and synthesizing the existing literature on these diets’ effects, focusing on growth performance, feed utilization, and gut microbiota. A systematic search of databases was conducted using Preferred Reporting Items for Systematic Reviews and Meta-Analyses (PRISMA) guidelines and a total of 57 studies were included in the qualitative analysis and 24 in the meta-analysis. The results indicated that soybean-based diets, at a 59.4% inclusion level improved the Specific Growth Rate (SGR) of tilapia with an effect size of -2.14 (95% CI: -2.92, -1.37; *p* < 0.00001; *I^2^*= 99%) and did not improve the feed conversion rate (FCR), as the effect size was 1.80 (95% CI: 0.72, 2.89; *p* = 0.001; *I^2^* = 100%). For seaweed-based diets, at a 15,9% inclusion level did not improve SGR, with an effect size of -0.74 (95% CI: -1.70, 0.22; *p* = 0.13; *I^2^* = 99%), and the FCR with an effect size of -0.70 (95% CI: -1.94, 0.54; *p* = 0.27; *I^2^* = 100%). Regarding the gut microbiota, was noted a lack of studies meeting the inclusion criteria for tilapia. However, findings from studies on other farmed fishes suggested that soybean and seaweed-based diets could have diverse effects on gut microbiota composition and promote the growth of beneficial microbiota. This study suggests that incorporating soybean-based diets at 59.4% inclusion can improve the SGR of tilapia. Seaweed-based diets, while not demonstrating improvement in the analyzed parameters with an inclusion level of 15.9%, have the potential to contribute to the sustainability of the aquaculture industry when incorporated at lower levels.

## Introduction

Tilapia are among the most important freshwater fish species worldwide and play a crucial role in global aquaculture production. As adaptable species to various environmental conditions, tilapia can be reared in a wide range of farming systems, including ponds, cages, and recirculation aquaculture systems [1,2,3]. The success of tilapia farming largely depends on the formulation of cost-effective and nutritionally balanced feed and an appropriate feeding management strategy that promotes optimal growth performance and feed utilization [4].

Traditional feed ingredients for tilapia include fishmeal and fish oil due to their high protein content balanced essential amino acids profile and omega-3 [5–8], but they are expensive and can contribute to the overfishing of wild fish stocks [9–11]. Therefore, there is a growing interest in exploring alternative protein sources for fish feed, such as soybean and seaweed. Soybean is a cost-effective, and widely available plant protein source [12,13], while seaweeds offer unique nutritional properties and are natural forage for fish [14]. Both have been investigated as partial or total replacements to improve the growth performance and feed efficiency of tilapia. Their inclusion in this study can provide a comprehensive understanding of their individual nutritional properties and highlight their potential as feed options for tilapia. However, there are challenges in including seaweed and soybeans in tilapia diets, such as mismatches in fatty acids and amino acids profile, leading to sub-optimal growth and health [15,16]. Additionally, soybean and seaweed contain antinutritional factors such as saponins, tannins, phytic acid, lectins, protease inhibitors, and amylase inhibitors [16–18], which may negatively affect nutrient digestibility, palatability, and overall feed performance.

Apart from the nutritional aspects, the gut microbiota plays a crucial role in the overall health and growth of fish [19–21] and can participate in modulating feeding behavior and efficiency [22,23]. Dietary interventions can influence the composition and activity of the gut microbiota, thereby influencing the host’s metabolism, immune system, and overall performance [24–27].

While individual studies have investigated the effects of soybean and seaweed-based diets on tilapia, a comprehensive understanding of their impact on growth performance, feed utilization, and gut microbiota is lacking. A systematic review and meta-analysis are essential to synthesize the available evidence and provide a comprehensive overview of the effects of these alternative feed ingredients on tilapia. Therefore, the present study evaluated and synthesized the existing literature on the effects of soybean and seaweed-based diets on growth performance, feed utilization, and gut microbiota of tilapia to achieve sustainability in aquaculture practices.

## Methods

### Study design

This study was conducted according to Preferred Reporting Items for Systematic Reviews and Meta-Analyses (PRISMA) guidelines (S1 Table). The protocol related to this systematic review was registered on Open Science Framework (<u>osf.io/zg72m</u>) with registration DOI: https://doi.org/10.17605/OSF.IO/ZG72M.

### Data source and search strategy

A systematic literature search was conducted using online databases including Google Scholar, PubMed, Wiley Online Library, and ScienceDirect, to identify relevant studies published between 2000 and 2022. The search terms were combined as follows: “seaweed” OR “macroalgae” NOT “microalgae” AND “growth performance” AND “feed utilization” AND “tilapia”; “soybean” NOT “soybean fermented” AND “growth performance” AND “feed utilization” AND “tilapia”; “seaweed” OR “macroalgae” NOT “microalgae” AND “gut microbiota” OR “intestinal microbiota” AND “tilapia” OR “other fish” NOT “human”; “soybean” NOT “soybean fermented” AND “gut microbiota” OR “intestinal microbiota” AND “tilapia” OR “other fish” NOT “human”. Additionally, a manual search of references cited in relevant articles was performed to identify additional studies that may have been missed in the database search. The search strategies and the respective number of studies from the four databases are detailed in the S2 Table.

### Inclusion and exclusion criteria

In this systematic review, studies published from January 2000 to December 2022 were included based on the following criteria: (1) articles written in English, (2) articles focusing on tilapia production; (3) studies utilizing various soybean-based diets such as soybean defatted, soybean meal, full-fat soybean, soybean milk, and soybean protein; (4) studies utilizing seaweed-based diets, including seaweed extract and various species of green, brown, red seaweed (S3 Table) to partially or totally replace fish meal/oil; (5) full-text available; (6) studies published in peer-reviewed journals; (7) studies reporting SGR and FCR with their respective standard error or standard deviation, and (8) articles examining the effect of soybean and seaweed-based diets on intestinal microbiota of other fish than tilapia. Excluded from consideration were: (1) studies not written in English; (2) studies not related to the use of soybean and seaweed as tilapia feed; (3) articles not published in peer-reviewed scientific journals; (4) articles not available in full-text format; (5) studies that used microorganisms to ferment soybean/seaweed; and (6) studies utilizing prebiotics in tilapia/fish production that were not derived from soybean or seaweed; (7) article that did not describe the composition of the gut microbiota; (8) studies lacking fish meal or fish oil as control group.

### Selection process

The screening of all papers was done in the Rayyan tool for systematic literature review (https://www.rayyan.ai/). After downloading the article in the databases (in RIS format), the files were imported to Rayyan to check for duplicates, screening the title, abstract, and full text by two independent reviewers (Leonildo dos Anjo Viagem-LAV and Jean Nepomuscen Hakizimana-JNH).

### Data extraction

Data were extracted from each included study using a standardized data extraction form created in Microsoft Excel 2013 (Microsoft Corporation, Washington, USA). This extraction process was carried out independently by two reviewers (LAV and JNH). The extracted data encompassed basic information such as author and publication year, experimental details including sample size and experiment duration, the inclusion level of the diets (both applied and recommended), and information about the different experimental groups, namely the soybean-based diet group, seaweed-based diet group, and control group (fish meal or fish oil). Parameters extracted for these groups included specific growth rate (SGR), and feed conversion ratio (FCR), along with their corresponding standard mean or standard error. Additionally, details regarding the types of soybean and seaweed species utilized in the experiments were recorded (S3 Table).

### Risk of bias in included studies

The study quality and risk of bias were evaluated using the Cochrane risk-of-bias tool in Cochrane Collaboration’s software Review Manager Version 5.4.1 (RevMan 5.4.1). This tool contains seven domains: (1) random sequence bias (selection bias), (2) allocation concealment (selection bias), (3) blinding of participants and personnel (performance bias), (4) blinding of outcome assessment (detection bias), (5) incomplete outcome data (attrition bias), (6) selective reporting (reporting bias), and (7) other biases [28]. This systematic review focused on five domains, as the blinding of participants and personnel (performance bias) and the blinding of outcome assessment (detection bias) are not applicable to aquaculture studies. In the domain of other bias was analyzed the information related to sample size and experiment duration.

### Meta-analysis

A meta-analysis was conducted to analyze the effect of soybean and seaweed on the SGR and FCR of tilapia. Effect sizes were calculated, and a random-effects model was employed. Publication bias was assessed using funnel plots of the effect of soybean on SGR and FCR (S1 Fig) and seaweed (S2 Fig). The heterogeneity of the included studies was assessed by the I-squared statistic test and was classified as low (<25%); moderate (25-50%) and high (>50%) [29]. The subgroup analysis and sensitivity were performed to explore possible causes of heterogeneity in the study. All statistical analyses were conducted in RevMan 5.4.1 and the variables were analyzed as continuous via standard mean difference with a 95% confidence interval (CI).

## Results

### Summary of included studies

Initially, a total of 17,991 studies were collected from the databases and 8 from other sources (Fig 1). After removing duplicates, 12,548 remained for screening. Based on eligibility criteria, 11,369 studies were excluded after titles and abstracts screening. The majority of the excluded studies focused on other animals than fish (n = 5,805) followed by those not addressing the topic of tilapia (n = 2,853) and seaweed or soybean (n = 2,351). Then, 1,179 full texts were assessed and 1,122 were excluded because most studies used probiotics to ferment soybean/seaweed (n = 910) and due to a lack of data on Specific Growth Rate or Feed Conversion Rate (n = 122). After full-text screening, 57 studies were included in the qualitative synthesis and 24 were included in the meta-analysis.

**Fig 1.**
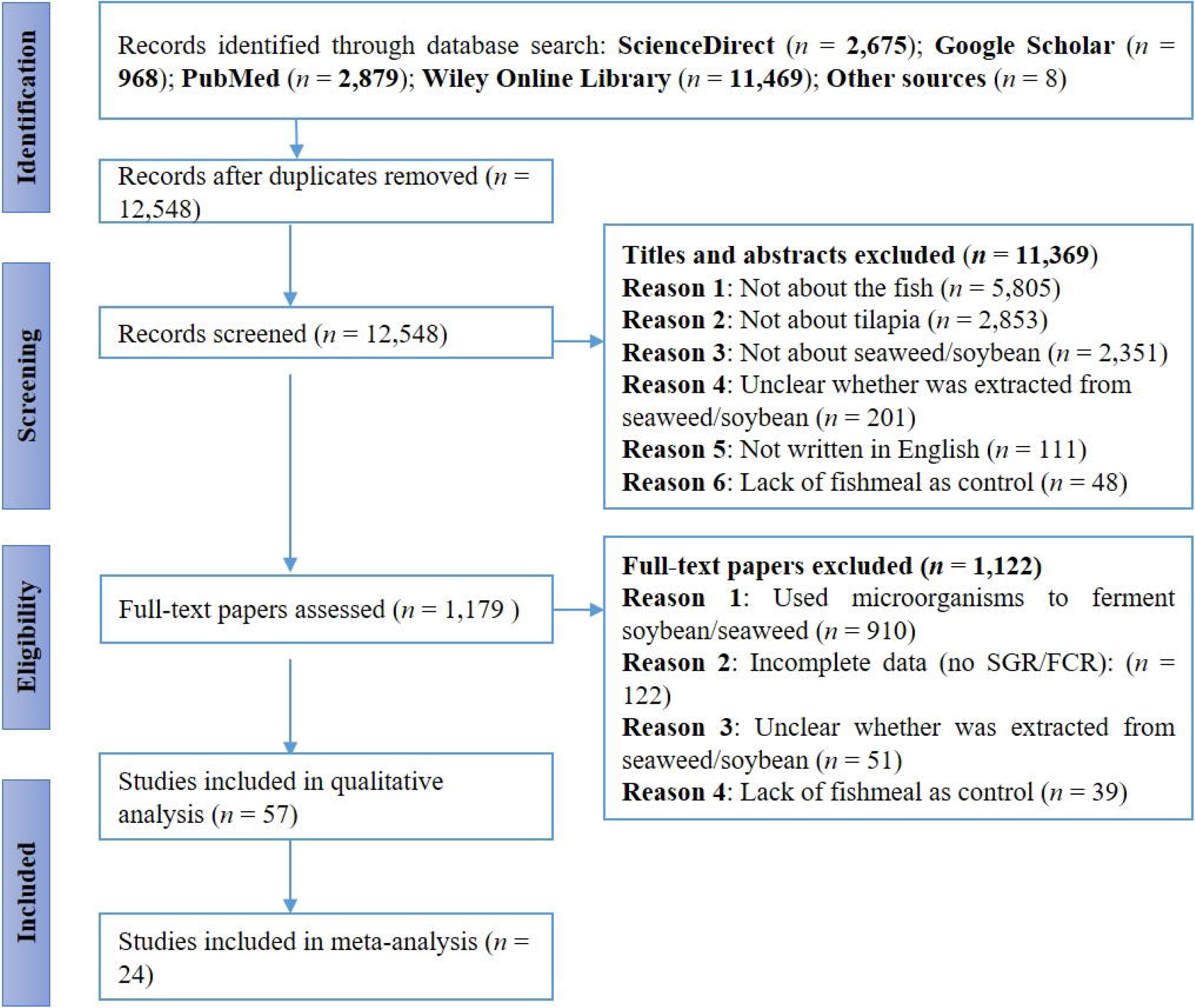
Preferred Reporting Items for Systematic Reviews and Meta-Analyses (PRISMA) flowchart of the study selection process. *n* represents the number of studies.

The included studies described that the fish were in good health and were purchased from companies that regularly conduct health inspections. These studies exhibited variations in experimental design, duration of the experiment, manner of processing soybeans, species of seaweed used in the experiment, and dietary inclusion level. In all the included papers, soybeans and seaweed were studied individually and in this review were analyzed separately.

### Quality and risk of bias assessment

The quality and risk of bias of included studies were assessed through the Cochrane criteria showing positive results for allocation concealment (selection bias), incomplete outcome data (attrition bias), selective reporting (reporting bias), and other bias (sample size and experiment duration) (Fig 2A and 2B). The evaluation of the studies revealed unclear risk bias for domain random sequence bias (selection bias) because most of the studies reported the random method but the sequence generation was not described. The studies Abdel-Warith et al., (2013), El-Saidy et al., (2010) and Sharda et al., (2017) (Fig 3A) and Stadtlander et al., (2012), Radwan et al., (2022) and Eissa et al., (2021) (Fig 3B) revealed high-risk bias for random sequence bias (selection bias). The study by Stadtlander et al., (2012) showed unclear bias for other bias because the sample size was not enough for this systematic review but the experiment duration was reported.

**Fig 2.**
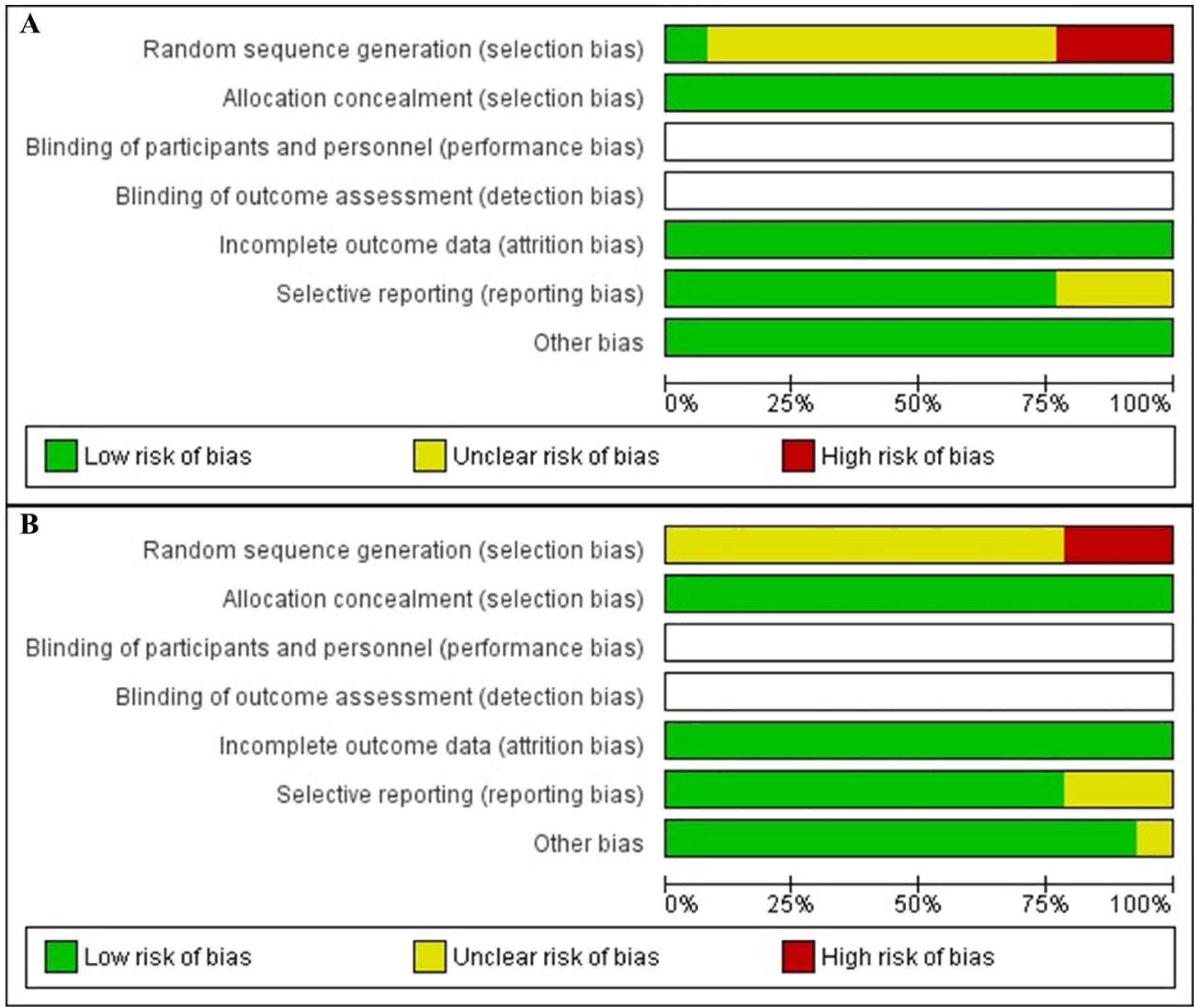
The risk of bias graph shows the review author’s judgment about each risk of bias item presented as percentages across all included studies on soybean (A) and seaweed (B). The quality and risk of bias was assessed using the Cochrane risk-of-bias tool in Cochrane Collaboration’s software RevMan 5.4.1

**Fig 3.**
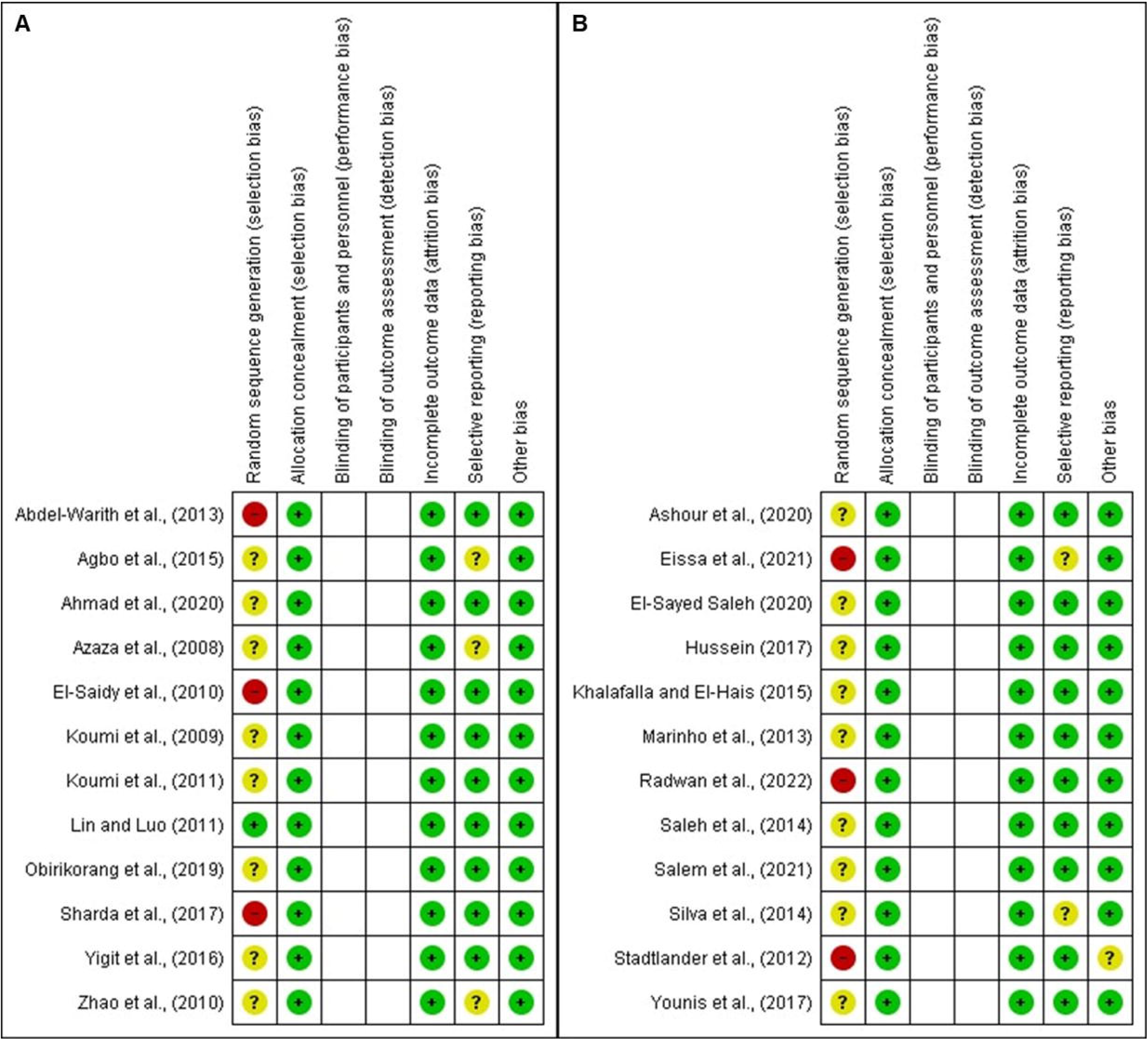
Risk of bias summary showing review author’s judgment about each risk of bias item for each included study on soybean (A) and seaweed (B). The quality and risk of bias was assessed using the Cochrane risk-of-bias tool in Cochrane Collaboration’s software RevMan 5.4.1

### Effect of soybean on growth performance and feed utilization

In this study, Specific Growth Rate (SGR) and Feed Conversion Rate (FCR) were used as parameters of growth performance and feed utilization, respectively. Among the studies included in the meta-analysis, twelve [11,16,30–39] described the effects of soybean on growth performance and feed utilization of tilapia (*Oreochromis niloticus* and *Sarotherodon melanotheron*) and involved 8,055 fish and 53 treatments. One article (Koumi et al., 2011) studied 2 species of tilapia, and the analysis was performed separately. The results showed that the random pooled effect size of SGR was -2.14 (95% CI: -2.92, -1.37; *p* < 0.00001; *I*^2^ = 99%) (Fig 4A) and for FCR was 1.80 (95% CI: 0.72, 2.89; *p* = 0.001; *I*^2^ = 100%) (Fig 4B). The average inclusion level of soybeans in the diets, calculated from the recommendation in each analyzed study (S3 Table), was found to be 59,4%. These findings suggest that including soybeans at this level can enhance the SGR but does not lead to an improvement in FCR.

**Fig 4.**
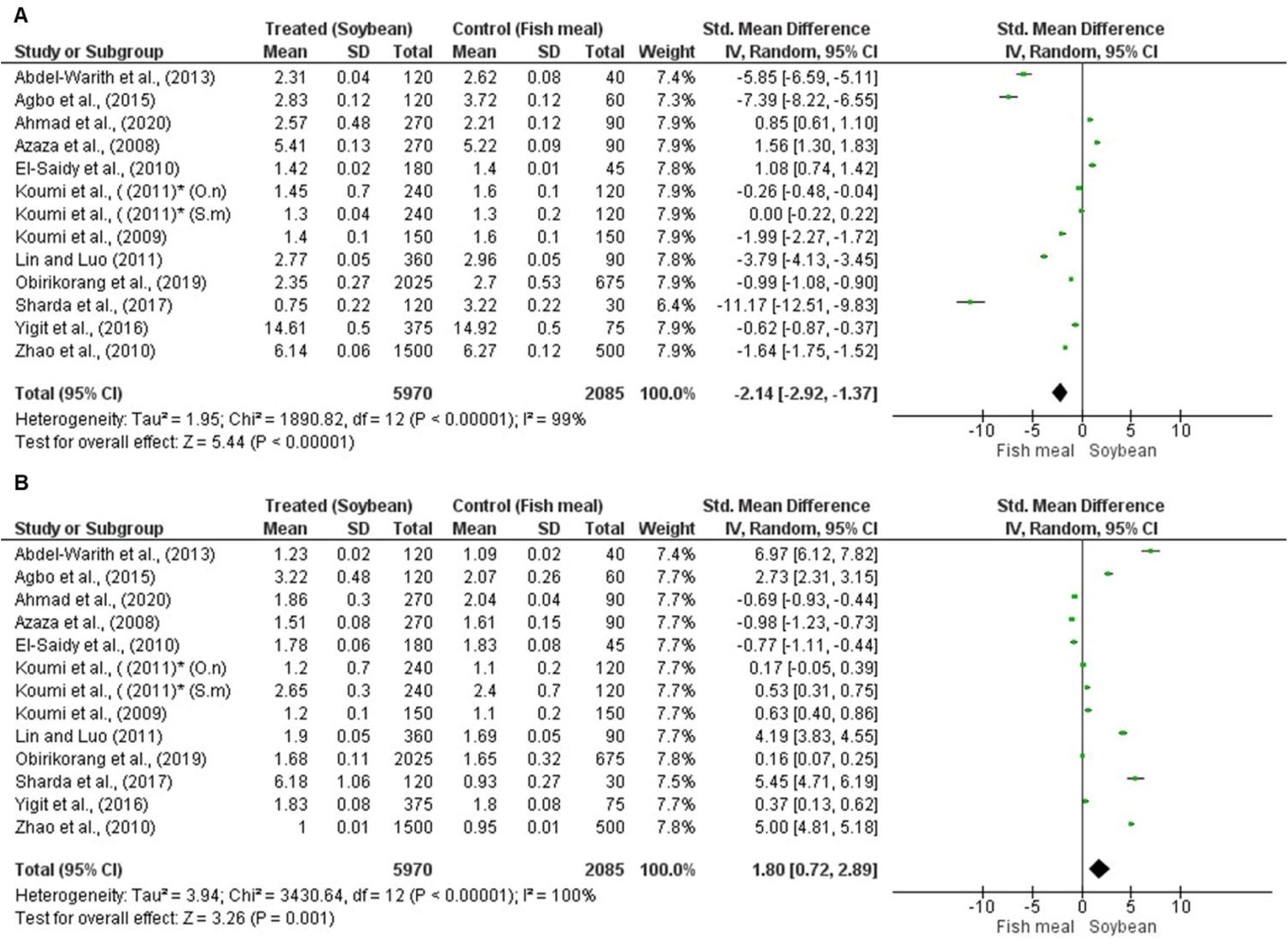
Forest plot showing the effect of soybean on specific growth rate (A) and feed conversion rate (B) of tilapia. The analysis was performed on RevMan 5.4.1, based on twelve studies and the effect size index is the standardized mean difference. The random-effects model was employed for this analysis and the studies in the analysis are assumed to be a random sample from a universe of potential studies. The variables were analyzed as continuous via standard mean difference with a 95% CI. ***O.n***: *Oreochromis niloticus*; ***S.m***: *Sarotherodon melanotheron*; **CI**: Confidence Interval; **Tau^2^**: tau-squared; **df**: difference; **Chi^2^**: chi-squared; ***P*:** *p*-value; **I^2^:** squared statistic test; **Z**: z-value; **SD** and **Std**: Standard Deviation; *same study but carried out on two species of tilapia (*O. niloticus and S. melanotheron*).

### Effect of seaweed on growth performance and feed utilization

Among the studies included in the meta-analysis, twelve [8,40–50] described the effects of seaweed on growth performance and feed utilization of *O. niloticus* and hybrid red tilapia (*O. niloticus* and *Oreochromis mossambicus*), involving 5,657 fish and 55 treatments of different species including rede seaweed (*Laurencia obtusa*, *Pterocladia capillacea*; *Gracillaria arcuata*, *Porphyra yezoensis*, *Gracilaria vermiculophylla,* and *Porphyra dioica*); green seaweed (*Ulva lactuca*, *Enteromorpha Flaxusa,* and *Ulva Fasciata*) and brown seaweed (*Taonia atomaria* and *Cystoseira myrica*) (S3 Table). Two papers (Silva et al., 2014 and Eissa et al., 2020) each studied three species of seaweed while one paper (Khalafalla and El-Hais, 2015) studied 2 species of seaweed, and the analysis was performed separately for each species. The results showed that the random pooled effect size of SGR was -0.74 (95% CI: -1.70, 0.22; *p* = 0.13; *I*^2^ = 99%) (Fig 5A), and for FCR was -0.70 (95% CI: -1.94, 0.54; *p* = 0.27; *I*^2^ = 100%) (Fig 5B). The inclusion level of seaweed was 15,9%. These results indicate that the inclusion level of seaweed at 15,9% did not improve the SGR and FCR of tilapia.

**Fig 5.**
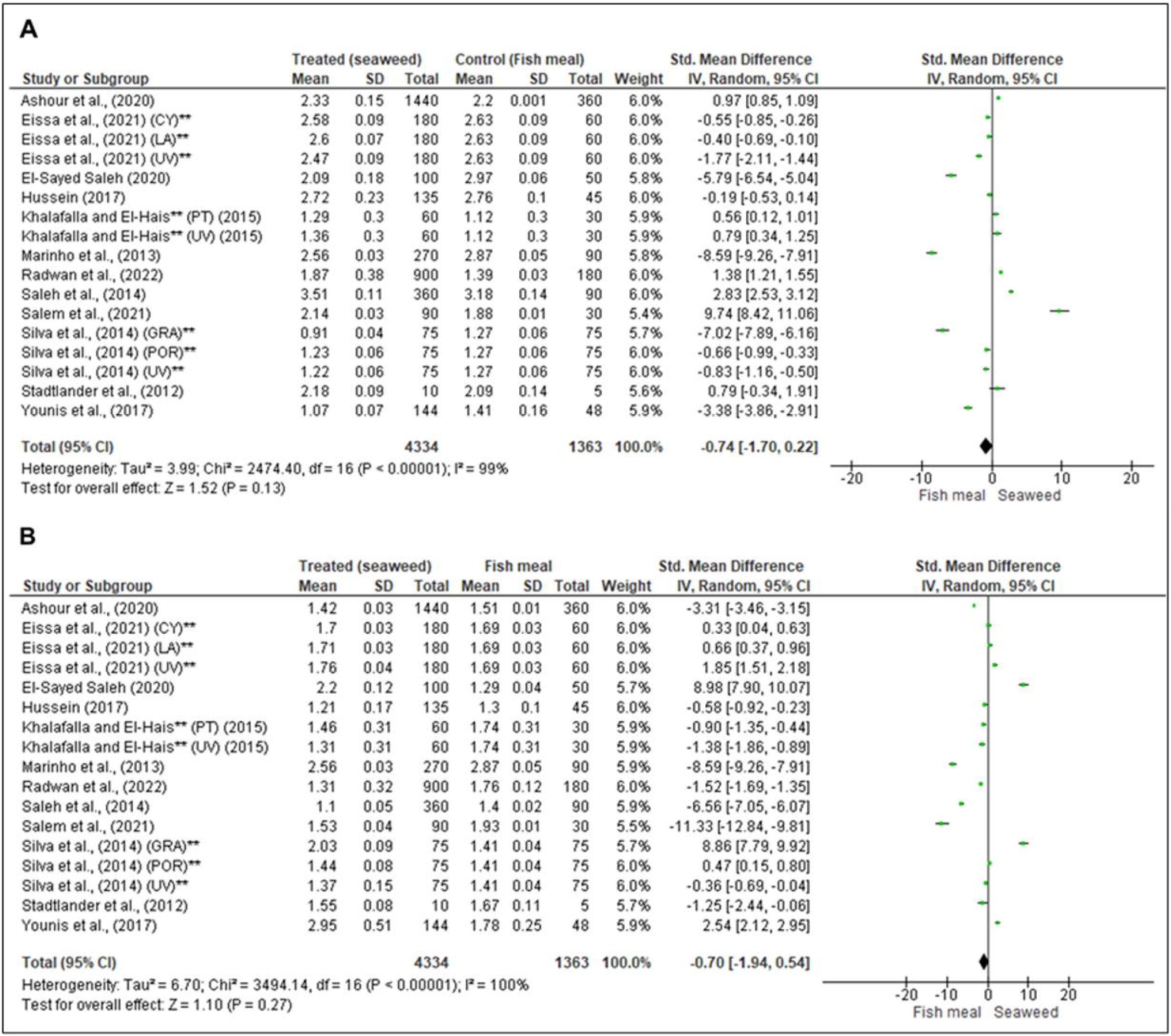
Forest plot showing the effect of seaweed on specific growth rate (A) and feed conversion rate (B) of tilapia. The analysis was performed on RevMan 5.4.1, based on seventeen studies and the effect size index is the standardized mean difference. The random-effects model was employed for this analysis and the studies in the analysis are assumed to be a random sample from a universe of potential studies. The variables were analyzed as continuous via standard mean difference with a 95% CI. **GRA**: *Gracillaria*; **UV**: *Ulva*; LA: *Laurencia*; POR: *Porphyra*; CY: *Cystoseira*; **CI**: Confidence Interval; **Tau^2^**: tau-squared; **df**: difference; **Chi^2^**: chi-squared; ***P*:** *p*-value; **I^2^:** squared statistic test; **Z**: z-value; **SD** and **Std**: Standard Deviation; **same study but carried out on different species of seaweed.

### Effect of soybean and seaweed on the gut microbiota of tilapia

The search conducted in the databases using the search strategies defined in this systematic review on tilapia was not able to identify studies that met the inclusion criteria on this specific topic. Most of the studies were done on other farmed fish (Table 1), showing that soybean and seaweed could have different effects on gut microbiota composition and diversity.

**Table 1.**
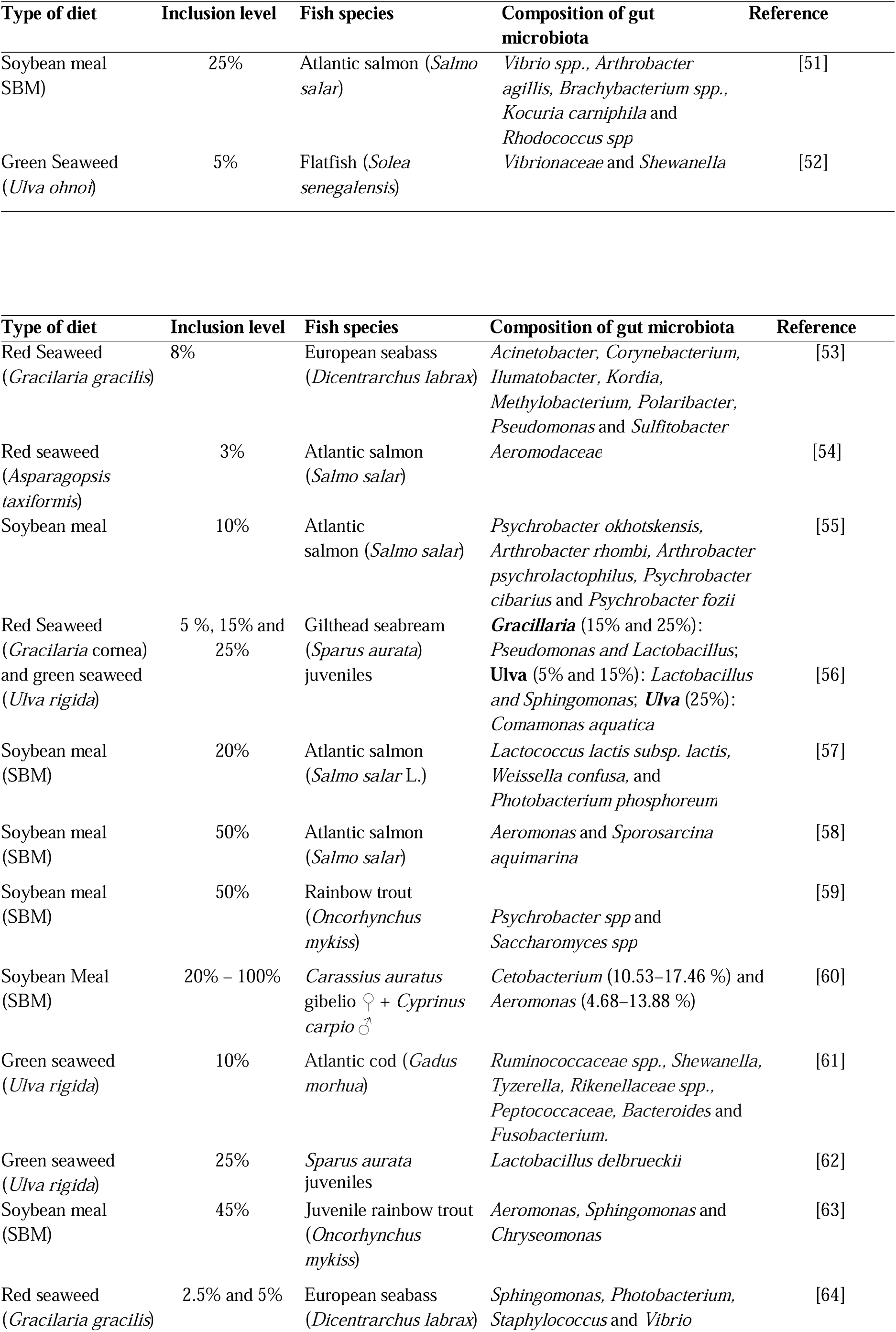
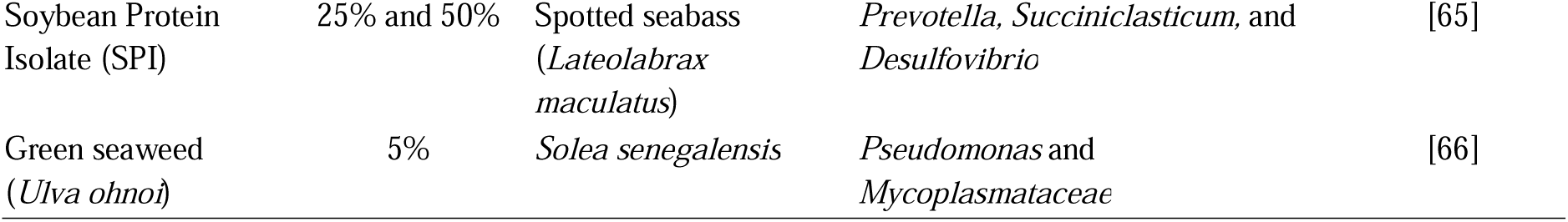
The most abundant gut microbiota in other fish than tilapia fed with soybean and seaweed-based diet.

## Discussion

### Effect of soybean and seaweed-based diet on SGR and FCR

The findings of this review suggest an enhancement in SGR among tilapia fed with soybean-based diets. This improvement could be attributed to soybean’s high protein content and relatively balanced amino acid profile [33,34]. The findings also reveal that soybean did not improve FCR, likely due to the presence of antinutritional factors such as protease inhibitors, lectins, phytic acid, saponins, phytoestrogens, antivitamins, and allergens [16,17]. These factors may compromise the palatability of soybeans and thus affect their effectiveness in improving FCR. The results from studies examining the impact of soybeans suggest that growth performance and feed utilization of tilapia tend to decline as the inclusion levels of soybeans increase, the ideal level of inclusion being up to 50% [31,35]. Similar studies conducted on other fish species such as *Liza haematocheila* [67]; *Heteropneustes fossilis* [68]; *Takifugu obscurus* [69]; *Clarias gariepinus* [70], *Solea aegyptiaca* [71]; *Sparus aurata* [72]; also indicate a decrease in growth parameters with higher levels of soybean inclusion level. However, determining the optimal inclusion level and type of soybean (e.g., defatted, fermented, full fat) requires further investigation since the studies included in the meta-analysis utilized different inclusion levels and types of soybeans. On the other hand, the results do not support the use of seaweed as an alternative protein source for tilapia, as it did not improve the SGR and FCR. However, it should be noted that the studies included in the meta-analysis used different species of seaweeds and different levels of inclusion.

Several studies have demonstrated that seaweed inclusion levels of up to 20% can effectively replace fishmeal in tilapia diets without compromising the growth performance and feed utilization [40,41,43,48,50,73]. Similar outcomes have been observed in other fish species as well [74–76].

The bioactive compounds present in soybeans and seaweed, such as Non-Starch Polysaccharides (NSPs) and Antinutritional Factors (ANFs) like saponins, lectins, and phytates, play significant roles in influencing fish digestibility and metabolic functions [77,78]. Excessive NSPs may cause gut inflammation and reduce energy availability, while ANFs can interfere with digestive enzymes and nutrient absorption [79]. However, proper processing methods such as heat and enzymatic treatment, bioprocessing, and fermentation of these diets can mitigate these effects [18,79,80]. Furthermore, phytochemicals like isoflavones in soybeans possess antioxidant properties, although their impact may vary among fish species [81]. Additionally, certain seaweed compounds, like polysaccharides may promote gut health, enhancing overall digestion and nutrient absorption [82,83]. However, the specific impact can vary depending on the type and amount of seaweed, as well as the fish species involved [15,84]. For instance, several studies have demonstrated that the green seaweed *Ulva spp*, when included at levels of up to 5%, can enhance the growth performance of certain carnivorous fish [85,86], and omnivorous fish like tilapia of up to 20% of inclusion [87]. In addition to soybeans and seaweed, plant-based diets such as sunflower, canola, cottonseed, Moringa, and *Azolla* meals have shown promise in feeding tilapia and other fish [88–92]. *Azolla*, noted for its exceptional amino acid profile, exhibits the highest nutritional index in tilapia feed [93]. In the data analysis, heterogeneity was observed among the included studies, potentially due to variations in experimental design, number of fish used in each study, type of soybean, and species of seaweed used across different studies. Subgroup analyses were performed to explore potential sources of heterogeneity, but the findings were inconclusive.

### Effect of soybean and seaweed-based diet on gut microbiota

In this review, it was not possible to find studies that describe the effect of soybean or seaweed on the gut microbiota of tilapia. However, studies in other fish species have shown diverse effects on the gut microbiota composition. For instance, beneficial bacteria like *Shewanella* were observed in Flatfish and Atlantic cod fed with *Ulva spp* [52,61], while *Lactobacillus spp* were found in Gilthead seabream fed with *Gracilaria cornea* and *Ulva rigida* [56,62] along with *Lactococcus spp* [57]. Additionally, *Psychrobacter spp, and Arthrobacter spp* were noted in fish fed with soybean meal [51,55,59]. These microorganisms are known to possess probiotic properties, which could potentially enhance nutrient absorption and disease resistance [94–97]. The seaweed and soybean dietary components contain bioactive compounds such as isoflavones, saponins, and oligosaccharides in soybean [98–100], and polysaccharides, polyphenols, phycobiliproteins, and phlorotannins in seaweed [101–103]. These compounds can exert prebiotic effects, influencing the composition of the gut microbiota [104]. Prebiotics are essentially non-digestible substances that, when consumed, foster beneficial changes in the gut microbiome, promoting the development of a more favorable microbial community [105]. This shift can have positive implications for various aspects of health, including digestive function, immune system modulation, and antimicrobial properties, inhibiting the growth of potentially harmful bacteria [106–108]. It is important to note that the specific effects of soybean and seaweed on fish gut microbiota can vary depending on various factors, such as the fish species, the inclusion levels of these ingredients, and the species of seaweed. Studies investigating the utilization of various woody forages such as *Moringa oleifera*, fermented *M. oleifera*, *Folium mori*, and fermented *F. mori* meals in tilapia farming have indicated significant variations in gut microbiota composition [109,110]. These findings suggest that diets incorporating soybean, seaweed, and other plant-based diets may have diverse effects on the diversity and composition of fish intestinal microbiota.

### Implications for tilapia production

The inclusion of soybean and seaweed-based diets into tilapia farming can contribute to the sustainability of the aquaculture industry by reducing reliance on fishmeal and fish oil. The improved SGR observed in tilapia fed with a soybean diet indicates the potential for cost-effectiveness and viability in tilapia production. Regarding seaweed-based diets, although they may not directly improve growth performance and feed utilization parameters in tilapia, they offer alternative feed options due to their rich content of bioactive compounds. These compounds can have beneficial effects on fish growth and health. Additionally, the modulation of gut microbiota by soybean and seaweed-based diets can improve tilapia health and nutrient utilization leading to more sustainable and cost-effective tilapia production.

### Limitations and future research directions

Despite the valuable insights gained from this study, limitations should be considered: (1) the included studies may have exhibited variations in experimental design, type of soybean and seaweed used, leading to heterogeneity in the data, (2) limitations for some outcomes, particularly effect of soybean and seaweed on gut microbiota of tilapia. For future research, it is suggested: (1) investigations into the optimal inclusion levels of soybean and seaweed-based ingredients and exploring the potential seaweed species for tilapia diets should be conducted to maximize growth performance and feed utilization; (2) long-term studies are needed to evaluate the potential impact of these diets on the gut microbiota and its implications for growth performance, nutrient utilization, and the health of tilapia and (3) investigate the effect of those diets in other species of tilapia.

## Conclusion

This systematic review and meta-analysis provide valuable insights into the effects of soybean and seaweed-based diets on tilapia production. The results of the meta-analysis indicated that the soybean inclusion level of 59.4% improved SGR but did not improve the FCR of tilapia. On the other hand, the seaweed-based diet did improve both SGR and FCR at an inclusion level of 15.9%. Based on the search strategies defined in this review, it was not possible to find studies that met the inclusion criteria for the gut microbiota of tilapia. However, studies on soybean and seaweed in other farmed fish have shown diverse effects on the gut microbiota composition and have the potential to enhance the growth of beneficial gut microbiota and improve fish health. This information is valuable for tilapia production as it offers an alternative dietary strategy for optimizing their growth.

## Support information

**S1 Table. Preferred Reporting Items for Systematic Reviews and Meta-Analyses (PRISMA) 2020 Checklist**

**S2 Table. Complete database search strategies and the number of articles accessed in each database**

**S3 Table. Data extraction from eligible studies on the effect of soybean and seaweed-based diets on tilapia production**

**S1 Fig. Publication bias represented by funnel plot of individual studies about the effect of soybean on SGR (A) and FCR (B)**

**S2 Fig. Publication bias represented by funnel plot of individual studies about the effect of seaweed on SGR (A) and FCR (B)**

## Author Contributions

**Conceptualization:** Leonildo dos Anjo Viagem

**Data curation:** Leonildo dos Anjo Viagem and Jean Nepomuscene Hakizimana

**Formal analysis:** Leonildo dos Anjo Viagem

**Methodology:** Leonildo dos Anjo Viagem

**Supervision:** Cyrus Rumisha, Brunno da Silva Cerozi and Gerald Misinzo

**Writing - original draft:** Leonildo dos Anjo Viagem

**Writing - review & editing:** Jean Nepomuscene Hakizimana, Cyrus Rumisha, Brunno da Silva Cerozi and Gerald Misinzo

## Data Availability

All data supporting the findings of this study are available within the paper and its supplementary information.

